# Changes in weak pair-wise correlations during running reshapes network state in the main olfactory bulb

**DOI:** 10.1101/2020.08.04.235382

**Authors:** Udaysankar Chockanathan, Emily J. W. Crosier, Spencer Waddle, Edward Lyman, Richard C. Gerkin, Krishnan Padmanabhan

## Abstract

Neural codes for sensory representations are thought to reside in a broader space defined by the patterns of spontaneous activity that occur when stimuli are not being presented. To understand the structure of this spontaneous activity in the olfactory system, we performed high-density recordings of population activity in the main olfactory bulb of awake mice. We found that spontaneous activity patterns of ensembles of mitral and tufted (M/T) cells in the main olfactory bulb changed dramatically during locomotion, including decreases in pairwise correlations between neurons and increases in the entropy of the population. Maximum entropy models of the ensemble activity revealed that pair-wise interactions were better at predicting patterns of activity when the animal was stationary than while running, suggesting that higher order (3rd, 4th order) interactions between neurons shape activity during locomotion. Taken together, we found that locomotion influenced the structure of spontaneous population activity at the earliest stages of olfactory processing, 1 synapse away from the sensory receptors in the nasal epithelium.

**New and Noteworthy:** The organization and structure of spontaneous population activity in the olfactory system places constraints of how odor information is represented. Using high-density electrophysiological recordings of mitral and tufted cells, we found that running increases the dimensionality of spontaneous activity, implicating higher-order interactions among neurons during locomotion. Behavior thus flexibly alters neuronal activity at the earliest stages of sensory processing.

## Introduction

Behavioral states, such as quiescence and wakefulness (Constantinople and Bruno 2011), locomotion (Dadarlat and Stryker 2017; Dipoppa et al. 2018) and arousal (Vinck et al. 2015) can have marked effects on patterns of neuronal activity throughout the brain. In sensory systems, neural activity can change in response to these behaviors, absent any sensory stimuli. This spontaneous activity often reflects the underlying organization of the circuit (Niell and Stryker 2010; Tsodyks et al. 1999), constraining what types of activity patterns are possible for encoding sensory stimuli (Luczak et al. 2009). These latent behavioral states consequently shape the variability of neuronal responses in primary sensory areas (Stringer et al. 2019)

As in the neocortex, neural activity in the main olfactory bulb is strongly influenced by behavioral state (Tsuno and Mori 2009). The principal neurons of the main olfactory bulb, the mitral and tufted cells (M/T) which receive direct input from the olfactory receptor neurons (ORNs), process these initial odor representations via connections with one another and with local inhibitory interneurons, and relay this information to downstream cortical areas (Wilson and Mainen 2006). Spontaneous firing rates in the principal M/T are lower under anesthesia as compared to waking (Rinberg et al. 2006a) and their activity can indicate non-olfactory information such as behavioral choice (Doucette and Restrepo 2008; Kay and Laurent 1999) or reward (Doucette et al. 2011). M/T cell firing can also be altered based on experience and training (Abraham et al. 2010; Kobayakawa et al. 2007; Smear et al. 2013). While the behavioral modulation of M/T activity is not surprising, given the link between olfaction, sniffing, and whisking (Carey and Wachowiak 2011; Kleinfeld et al. 2014; Moore et al. 2013; Verhagen et al. 2007), the fact that it occurs in principal cells one synapse away from the sensory periphery (Murthy 2011; Wilson and Mainen 2006) suggests that the impact of behavior on neuronal coding may occur earlier in olfactory processing (Tsuno et al. 2008) as compared to other sensory systems in mammals. Centrifugal projections into the bulb (Padmanabhan et al. 2016; Price and Powell 1970, 1971; Shipley and Adamek 1984) from a diversity of areas including in the hippocampus (Davis and Macrides 1981; Padmanabhan et al. 2018), point to a density of anatomical connections that link olfactory processing in the bulb to a complex array of behaviors (Gourevitch et al. 2010) including running. While a number of recent studies suggest that locomotion and running exert substantial influence on population activity in sensory neocortices (Dadarlat and Stryker 2017; Dipoppa et al. 2018; Erisken et al. 2014; Stringer et al. 2019), it is less clear how locomotion, which is essential for a number of olfactory behaviors including foraging (Gire et al. 2016), and tracking (Khan et al. 2012), influences the activity of ensembles of neurons in the earliest stages of olfactory coding.

To address this, we recorded simultaneously from 100s of M/T cells in main olfactory bulb (MOB) while animals ran on a cylindrical wheel. Not only did running increase firing rates in individual neurons, running also change the structure of activity across populations of cells, including increased the entropy of the ensemble activity. An Ising maximum entropy model of M/T cell firing patterns uncovered that small changes in the pair-wise interactions between neurons during locomotion had large effects on the global structure of population activity when the animal was running. During locomotion, decreases in pair-wise interactions reduced the predictive power of 2^nd^ order maxent models, opening up the possibility that higher order structure in the shapes neural population activity.

## Materials and Methods

### Animals

All protocols and procedures were approved by the University Committee on Animal Resources (UCAR) at the University of Rochester and were performed in accordance with the guidelines of the Institutional Animal Care and Use Committee (IACUC) at the University of Rochester. 3 female and 2 male C57Bl6/J mice, age 3-6 months, were used in this study.

### Surgery

Mice were anesthetized with an inhaled 1-2% isoflurane mixture. The scalp was removed and a 3D printed headframe was attached to the dorsal skull surface with veterinary adhesive (Vetbond, The 3M Company, Maplewood, MN, USA) and dental cement (Ortho-Jet Powder and Jet Liquid, Lang Dental Mfg. Co., Wheeling, IL, USA). A metal ground screw was also implanted into the skull. Following surgery, mice were monitored until they recovered from anesthesia. Post-operative analgesia was provided in accordance with approved protocols and animal weight was monitored during the week.

### Run-wheel training

For a 5-day period following headframe implantation, mice were placed on a non-motorized running wheel for 1 hour daily. During these sessions, electrophysiological recordings were not performed. The purpose of these sessions was to acclimate the mice to running on a wheel while head-fixed. We previously found that following 5-days of exposure, both male and female mice habituate to the run wheel based on their running behavior, after which electrophysiology was performed (Warner and Padmanabhan 2020).

### Electrophysiology

Mice were anesthetized with 1-2% inhaled isoflurane and a craniotomy was performed over the right main olfactory bulb using stereotactic coordinates (5 mm Rostral, 1 mm lateral, and 1-1.5 mm ventral of bregma). Immediately following the craniotomy, mice were transferred to the running wheel, where they were head-fixed and allowed to recover from anesthesia. A 128-channel nanofabricated silicon electrode array (Du et al. 2011) was vertically lowered into the MOB (**Fig. 1A,B**). Extracellular voltage recordings were collected at 30kHz in the 0.1-3500Hz frequency band (**Fig. 1C**). For four of the mice, one recording session was performed. For one mouse, a second recording session in a different region of the MOB was performed. Simultaneously, the running velocity of the mouse was also recorded via a rotational encoder attached to the wheel. All recordings were performed while mice were awake and behaving on the non-motorized voluntary running wheel. The recording rig was enclosed in a box to minimize interference from ambient odor, light, sound, and electromagnetic noise.

**Figure 1:**
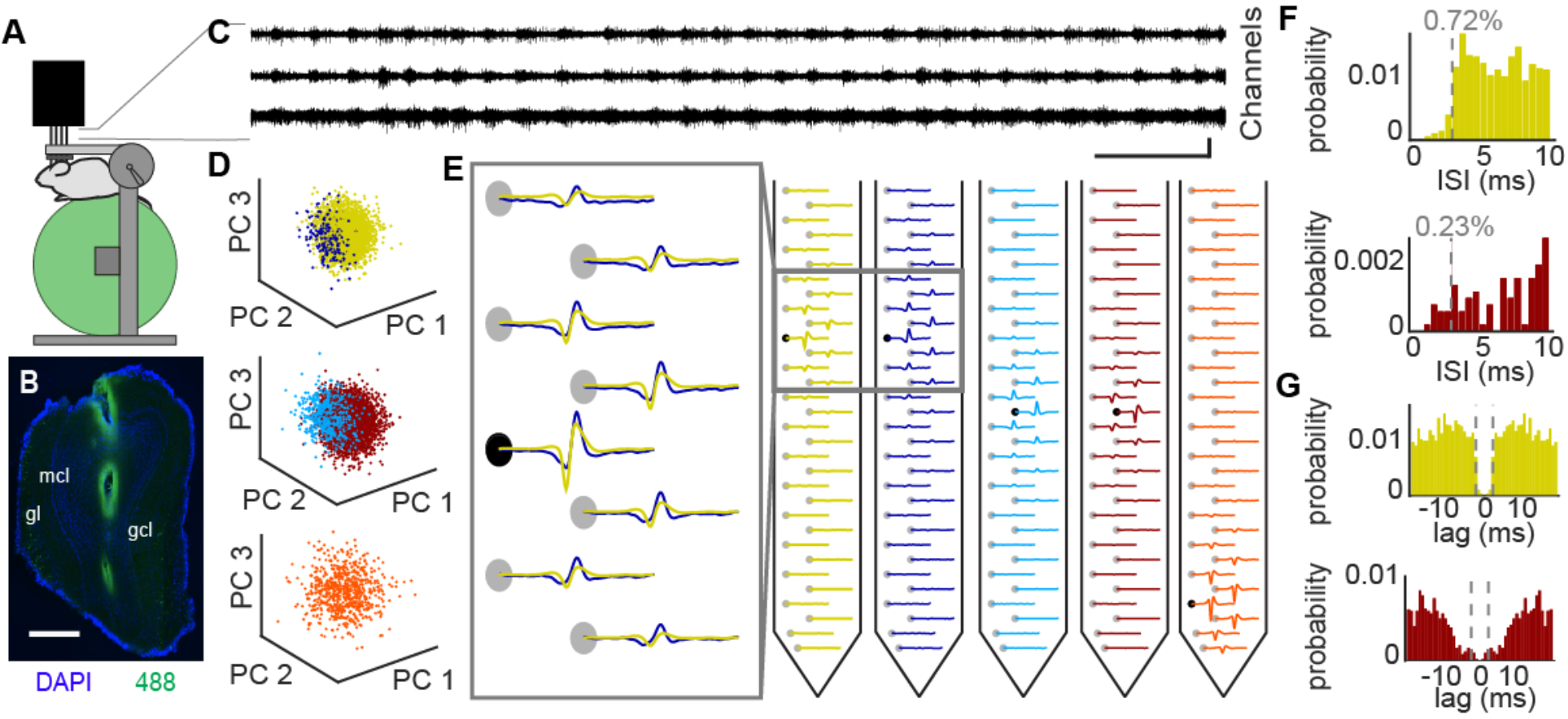
Electrophysiological monitoring of large ensembles of mitral/tufted cells in a head-fixed animal on a run wheel. (A) Schematic of recording population activity from Mitral/tufted cells in the main olfactory bulb of awake mice affixed to a run-wheel. (B) Coronal section of MOB showing recording location (labeled by cholera toxin B subunit conjugated to Alexa 488). Scale bar = 100 μm. (C) Bandpass (500-3500 Hz) voltage trace from 3 channels of 128 channels show spiking activity across multiple contacts. (D) PCA projections of spike waveform clusters used for identifying single units. Each color represents a waveform from E, each point is a single spike. (E, left) Spike waveforms of two units across 8 channels from a single 32 channel shank. (E, right) Waveforms across all 32 channels for 5 units show that spiking activity from a single cell can be found across 8-12 contacts. (F) Inter-spike interval histogram for two units in D and E. (F) Auto-correlation of spike counts from two units in D and E.

### Spike sorting

Spike sorting was performed as previously described (Chockanathan et. al. 2020). Briefly, putative action potentials, or spikes, were identified as intervals during which the band-passed (500-3500 Hz) voltage magnitude exceeded 6 standard deviations from baseline. Periods containing voltage artifacts were manually identified and eliminated.

Spikes were assigned to units, or putative single neurons, by concatenating all the spikes on a channel with the signal from 8 neighboring channels and projecting them onto a space of principal components (PC) that explained 80% of the variance. Clusters were identified automatically using a mixture of Gaussians model (Lewicki 1998) and final manual curation was performed using criterion including the number of total spikes for that unit (>50), mean waveform shape, spike autocorrelogram, interspike interval, and waveform similarity across units.

### Running behavior analysis

The motion of the run-wheel was controlled by the mouse, which could ambulate in forward or reverse directions or remain stationary. Wheel movement was converted to a velocity by integrating with a 150ms time bin in 10ms steps. Bins in which the velocity exceeded 1cm/s were classified as running and all other bins were classified as stationary.

### Correlations

A sliding window was applied to the spike rasters to convert the binary spike pattern to a continuous firing rate trace, which was then converted to a z-score. The correlation coefficient between the running velocity and the activity of every unit in each animal was calculated. Null distributions of correlations were calculated by temporally shuffling spike times and generating instantaneous firing rate time-series from the shuffled spike trains.

### Entropy

The spike train of each unit was binarized using 10ms non-overlapping bins. If one or more spikes from a given unit was present in a bin, the bin was assigned a value of 1. If no spikes were present in that bin, it was assigned a value of 0. This resulted in a binary matrix, in which each column describes the state of every neuron in the recorded population at a given time (Pryluk et al. 2019; de Ruyter van Steveninck et al. 1997; Schneidman et al. 2006). The probability distribution of these population states (entropy) was calculated as:

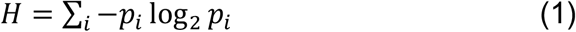

where *H* is the entropy and *p_i_* is the probability of the *i*^th^ pattern. For entropy analyses, the overall recorded population of neurons from each animal was subsampled 500 times. The number of neurons included in the subsample was varied between n = 3 neurons and 25 neurons. Entropy for running and stationary epochs was conditioned on the run velocity being greater than 1cm/sec

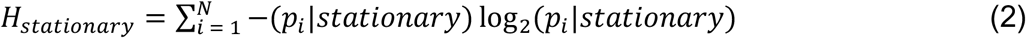

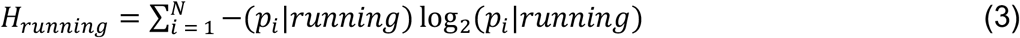

### Maximum entropy models

The *maxent_toolbox* software (Maoz and Schneidman 2017) was used to fit the maximum entropy models (Ohiorhenuan et al. 2010; Schneidman et al. 2006). First, the local field terms *h_i_* and pairwise interaction terms *J_ij_* were computed from the binned spike trains. The resulting model was then used to predict the probability of each pattern:

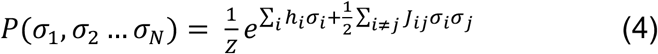

where *σ_i_* denotes the binary state of each neuron and Z denotes the partition function, which normalizes the pattern probability distribution. The pattern probabilities predicted by the model were then compared to those observed in the data using the Kullback-Liebler divergence (KLD). The data binarization process used for the maximum entropy models is the same as that used for the entropy calculations.

## Results

To study the effect of locomotion on spontaneous neuronal population activity, high-density 128 channel arrays were targeted to the MOB in head-fixed C57Bl6/J mice (n = 3 females, 2 males, age 3-6 months) trained to run on a non-motorized wheel (**Fig. 1A,B**). Action potentials in the extracellular recordings (**Fig. 1C**) were identified, and single-unit activity was clustered using a mixture-of-Gaussian models (Lewicki 1998) (**Fig. 1D-E**). Representative mean waveforms from putative M/T cells across all the channels in the electrode shank (**Fig. 1E**) illustrate the waveform diversity of units, with inter-spike interval (ISI) distributions (**Fig. 1F**) and autocorrelation functions (**Fig. 1G**) reflecting the quality of spike-sorting (see methods). On average, 113 ± 46 well-isolated units were identified per recording session (**Fig. 2A**).

**Figure 2:**
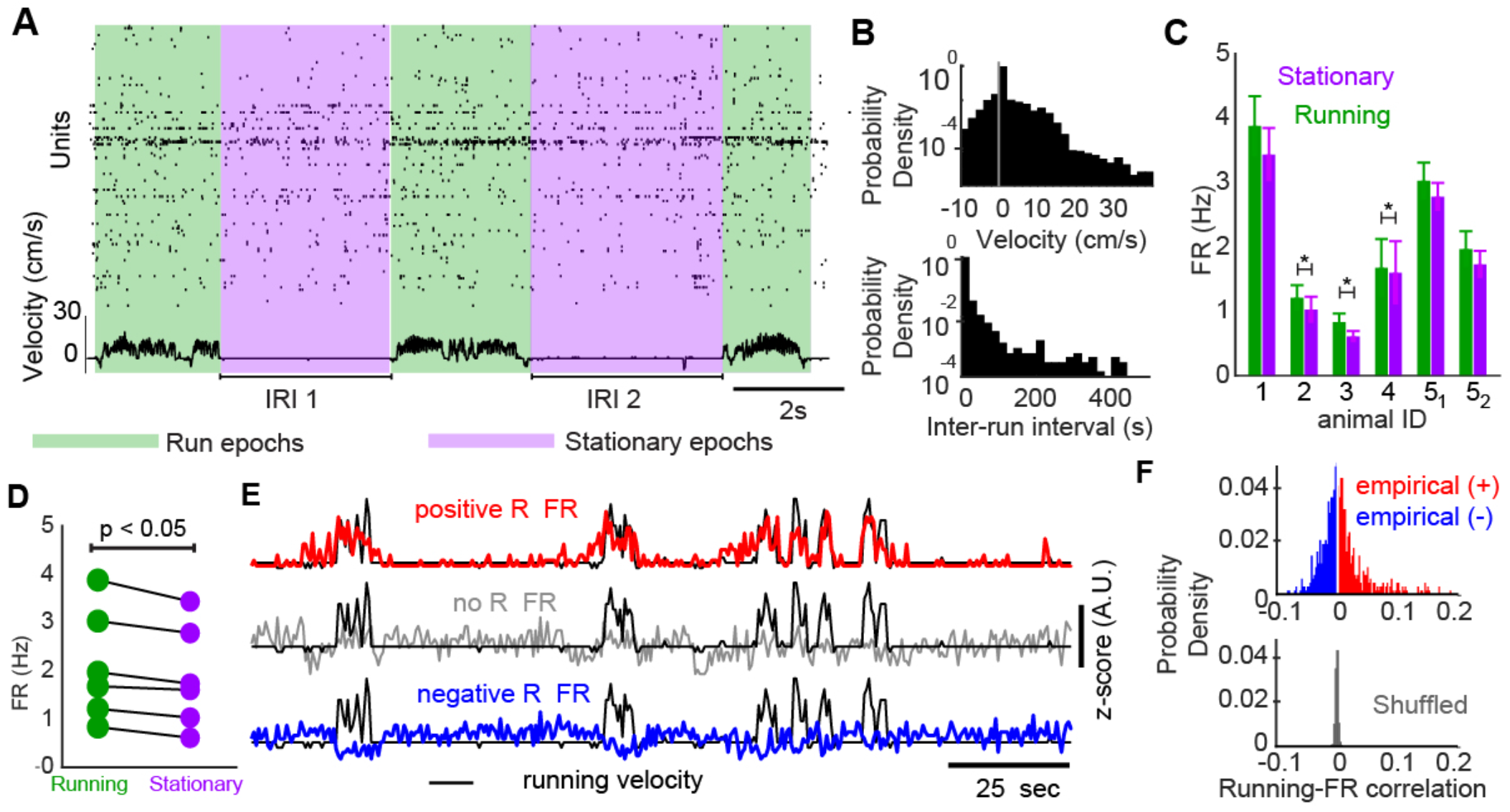
M/T cell firing was modulated by locomotion. (A) Example trace of ensemble of recorded units while the animal was running (green) versus when it was stationary (purple). (B, top) Histogram of run velocity across all animals. (B, bottom) Histogram of inter-run intervals. (C) Mean firing rate across units for 6 recording sites across 5 animals. Firing rates were significantly higher in 3 animals when the animal ran. (D) Firing rates were significantly higher on average during running across all recording sites from all animals (N = 6). (E) Example traces of unit firing rate and running velocity for a correlated unit (red), uncorrelated unit (gray), and negatively correlated unit (blue). (F) Histogram of correlations between unit firing rate and run velocity (top, positive correlated units = red, negatively correlated units = blue, bottom, shuffled data = gray).

Throughout the recordings, intervals of when the mouse remained stationary were interspersed with epochs of running (inter-run interval as defined as the epoch between the end of one run interval and the beginning of another = 6.2 ± 25.6s, **Fig. 2B**). In all animals, neuronal activity was dynamic both across and within stationary and running epochs (**Fig. 2A**). At the single-animal level, mean firing rates during running were significantly larger than those during stationary epochs in three animals (Bonferroni-corrected p < 0.05, two-sided Wilcoxon signed-rank test, **Fig. 2C**). At the group level, overall firing rates were increased with running (FR_stationary_ = 1.86 ± 1.06Hz, FR_running_ = 2.10 ± 1.15Hz, p < 0.05, two-sided Wilcoxon signed-rank test, N = 6, **Fig. 2D**). Although average firing rates across the population increased with running, individual neurons exhibited diverse changes in their activity patterns including both increases and decreases in firing during locomotion. For example, a plot of the run velocity of the animal and the firing rates for three example M/T cells showed units that were negatively correlated, uncorrelated, or positively correlated with running velocity (**Fig. 2E**). The range of correlations observed was significantly different than chance, as determined using temporally shuffled spike trains (p < 10^-6^, two-sample F-test, **Fig. 2F**).

As odor-encoding in the bulb is done not by single units, but relies on the activity of ensembles of mitral and tufted cells from both the same (Dhawale et al. 2010) and different glomeruli (Gerkin et al. 2013), we wished to examine how locomotion sculpted the *collective* activity of neuronal populations in the MOB. First we visualized the population activity of M/T ensembles in a low dimensional space defined by the first three principle components calculated from the covariance of firing patterns across the entire recording duration for two example animals (**Fig. 3A1-2**). In these low-dimensional embeddings, ensemble activity when the animal was running was differentially distributed as compared to periods when the animal was stationary (**Fig. 3B1-2**). As a result, not only the structure of single neurons, but the activity of populations of cells was different during locomotion. These visualizations suggested that this collective neural activity resided in different states in a high-dimensional space depending on the animal’s locomotor behavior. However, it is difficult to compare changes in neural activity during locomotion across animals as differences in the dimensionality for each animal depend on analysis method. We instead turned to a measure of statistical dispersion of firing patterns in M/T cells, entropy, that quantifies the diversity of states across ensembles when the animal is running versus when it is stationary. To do this, we binarized spike trains from subsampled ensembles of M/T cells into patterns of 1s and 0s and calculated the entropy of the activity when the animal was stationary versus when it was running (**Fig. 4A**). For 500 resamples of 8-unit subpopulations in a single animal, entropy (denoted-H) was greater when running as compared to when the animal was stationary (H_stationary_ = 77.4+28 bits/sec, H_running_ =85.7+35 bits/sec, p < 0.05, two-sided Wilcoxon signed-rank test, **Fig. 4B**). The increase in entropy was also significant across the all recording sessions in animals (H_stationary_ = 82.7+11 bits/sec, H_running_ = 91.8+13 bits/sec, p < 0.005, two-sided Wilcoxon signed-rank test, N = 6, **Fig. 4C** inset). Furthermore, the entropy was significantly larger with running over a range of subpopulation sizes from 2 to 25 neurons (p < 0.005 for all populations from 2-25 neurons, two-sided Wilcoxon signed-rank test with Bonferroni correction, **Fig. 4C**). Additionally, the entropy was larger during locomotion across a variety of additional bin sizes (50 ms, 100 ms, data not shown), meaning the number of patterns of spontaneous activity when the animal was locomoting were larger.

**Figure 3:**
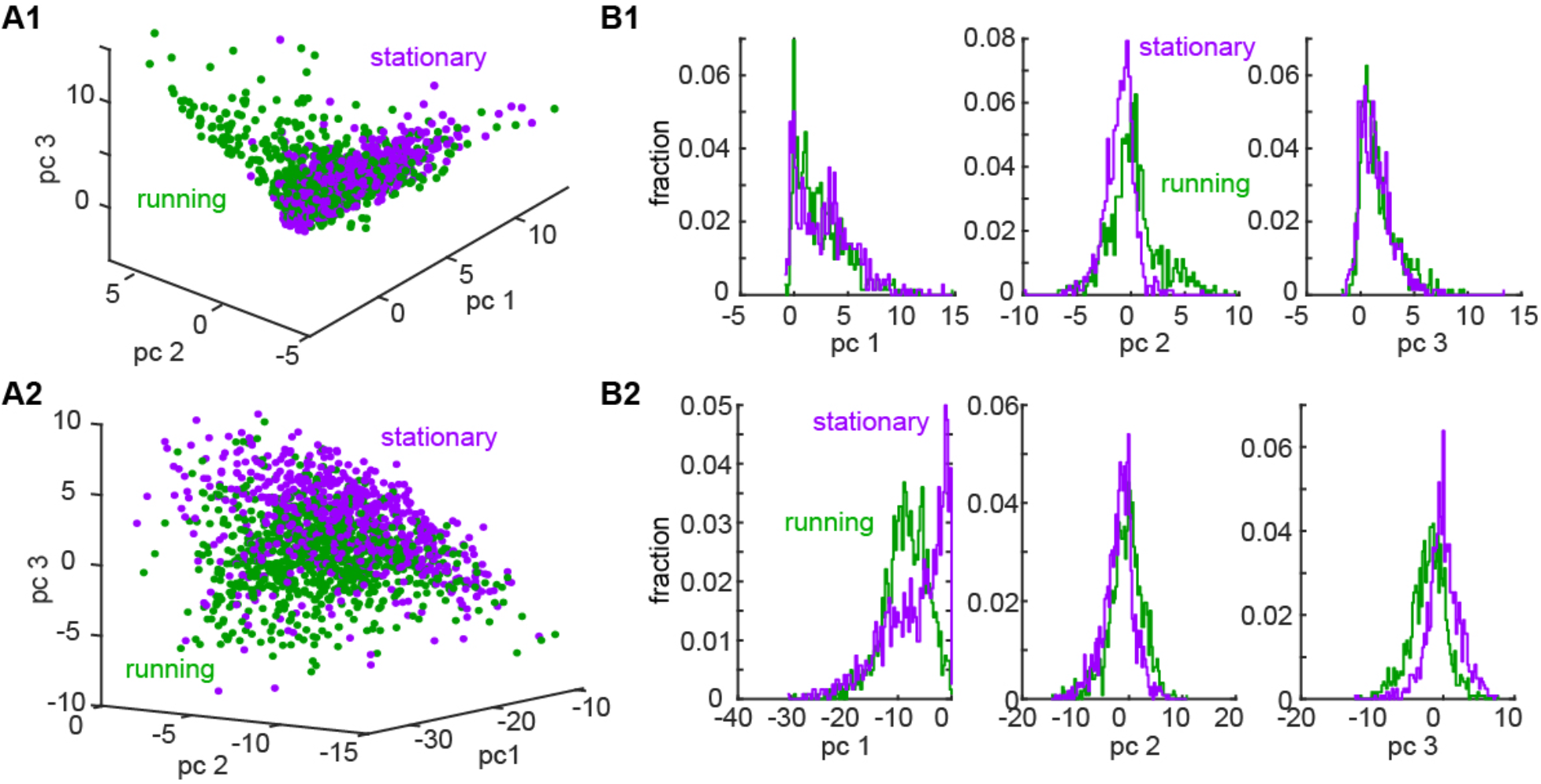
Ensemble activity was differentially distributed in a state space during running vs stationary epochs. (A) Two representative examples of ensemble activity from > 100 simultaneously recorded units projected onto a low dimensional space when the animal was running (green) versus when it was stationary (purple). Each dot corresponds to a single time. (B) Histograms of activity during running versus when the animals were stationary in each of the three dimensions in A.

**Figure 4:**
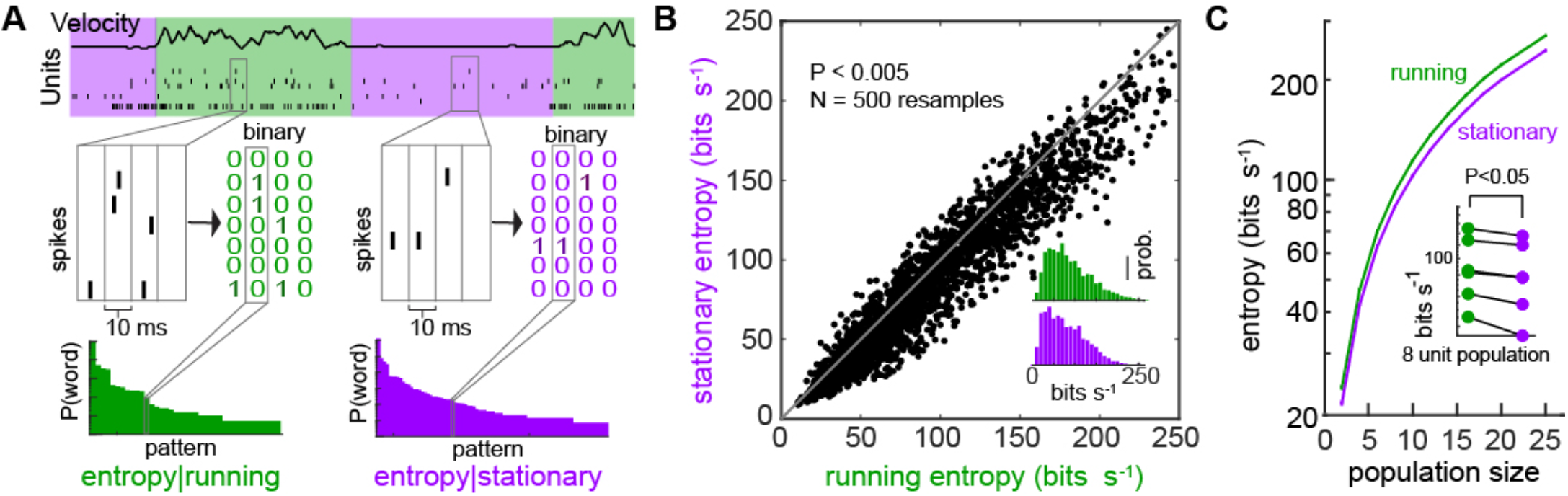
Entropy was greater when animals were running as compared to when they were stationary. (A) Schematic describing how ensemble activity was converted into binary patterns. Spikes were binned in 10 ms windows and converted to patterns of 1s and 0s. (B) Entropy for 500 subsamples of an 8 neuron population in a single animal during running versus stationary epochs. (B, insert) Histogram of entropy for running versus stationary epochs. (C) Entropy for different subpopulations sizes from 2 to 25 units (500 samples/animal/subpopulation size) shows that entropy was significantly higher when the animal was running as compared to when it was stationary. (C, insert) Mean entropy was lower when stationary as compared to running at the animal level.

We next wished to identify what features of the ensemble activity gave rise to the changes in the entropy, and what these might reveal about how the structure of population activity is shaped by behavior. To do this, we constructed maximum-entropy models of M/T cell activity, which incorporate as few *a priori* assumptions about population interactions as possible (Chockanathan et al. 2020; Meshulam et al. 2017; Schneidman et al. 2006). These models contain a local field term (**h_i_**) for each neuron as well as a coupling term (**J_ij_**) for each pair of neurons. The local field term **h_i_**, is a measure of how likely the cell is to spike, the more negative, the more silent the firing of the neuron. The **J_ij_** terms captured the interactions, the functional coupling across all pairs of cells. Both terms were calculated directly from the population activity when the animal was stationary versus when it was running (**Fig. 5A**), effectively identifying how the baseline firing and pair-wise interactions between neurons were altered when the animal ran. Using these terms for pairs of neurons, we then predicted the frequency of patterns across the population that should occur based on pair-wise interactions and compared the predicted patterns to those observed in the data (**Fig. 5B** representative 8-unit subsample from a single animal). The closer to the unity line (black), the better the model was at predicting patterns of activity. In this example, predictions of patterns from the model during running were worse than those during stationary epochs (**Fig. 5C**). The difference in prediction between the model and the data was quantified using the Kullbeck-Liebler divergence (KLD). For example, when comparing the fits at the group level for an 11-unit population, the KLD was increased for running epochs as compared to stationary epochs, indicating that the fit was worse when the animals ran (KLD_stationary_ = 0.0012 ± 0.0008, KLD_running_ = 0.0040 ± 0.0015, p < 0.05, two-sided Wilcoxon signed-rank test, N = 6, **Fig. 5D** inset). Across a range of subpopulation sizes (3 to 19 units), the pair-wise maximum entropy model was worse at predicting the global patterns of activity when the animal was running as compared to when it was stationary (p < 0.05, two-sided Wilcoxon signed-rank test with Bonferroni correction, **Fig. 5D**).

**Figure 5:**
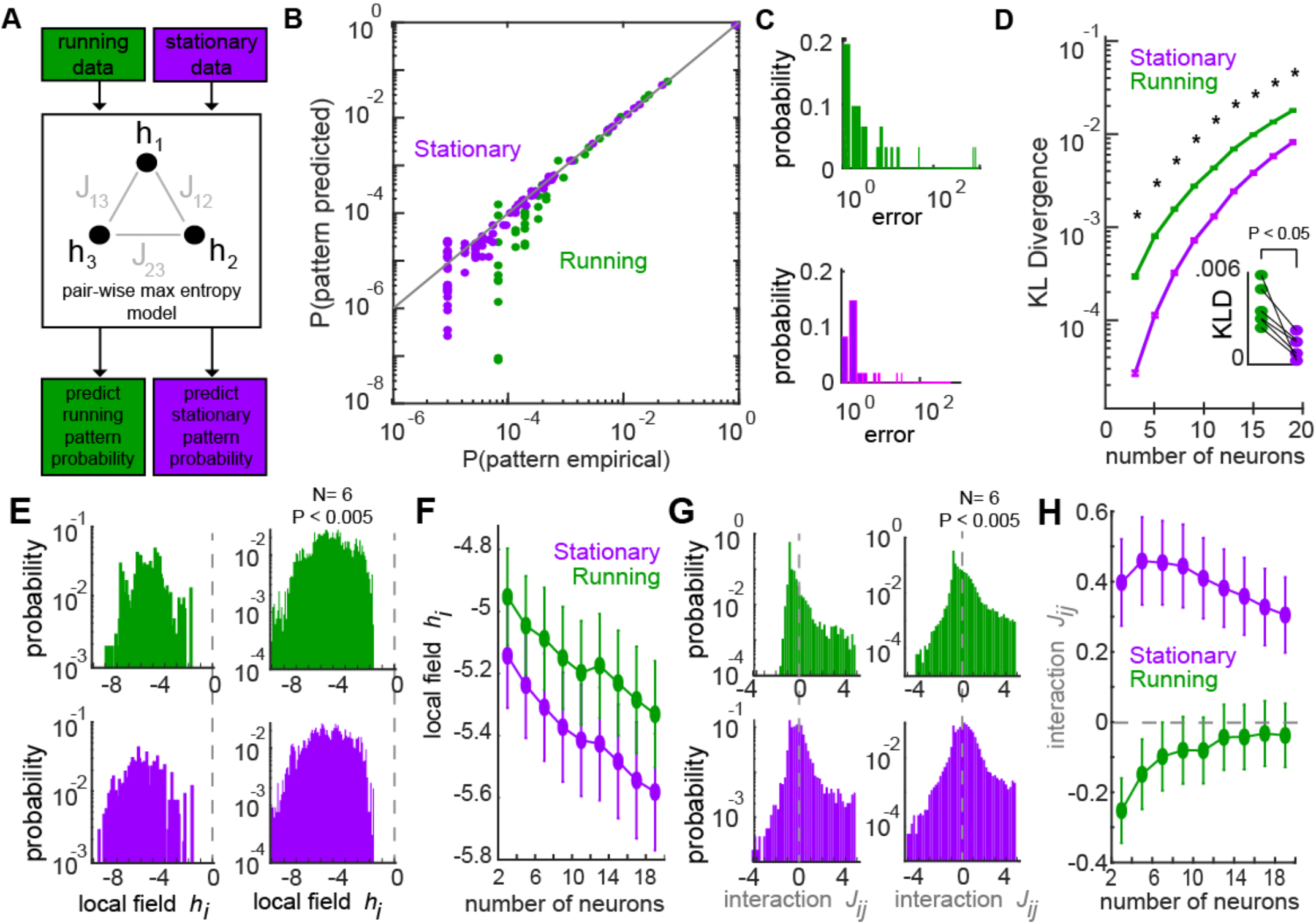
Pair-wise maximum entropy models were better at fitting M/T cell ensemble activity when the animal was stationary as compared to when it was running. (A) Schematic of method for calculating pair-wise maximum entropy models. (B) For an example 8-unit population, the rate occurrence of each firing pattern from the pair-wise maximum entropy model was plotted against the observed data for running (green) and stationary (purple) epochs. (C) Histograms of prediction error of rate of occurrence of the firing patterns in B showed that error was higher when the animal was running. (D) KL divergence, which measures the goodness of pair-wise maximum entropy model prediction for different subpopulations sizes from 3 to 19 units during running versus stationary epochs. (D, insert) Mean KLD for each recording session during running versus stationary epochs. (E, left) Histogram of local field term of pair-wise maximum entropy model from a single animal across multiple resamples of an 8 unit subpopulation. (E, right) Histogram of local field term of pair-wise maximum entropy model from all recording sessions. (F) Mean local field term for different subpopulation sizes. (G, left) Histogram of interaction terms of pair-wise maximum entropy model from a single animal across multiple resamples of a 8 unit subpopulation. (G, right) Histogram of local field term of pair-wise maximum entropy model from all recording sessions. (H) Mean interaction term for different subpopulation sizes.

To understand what the maximum entropy models revealed about ensemble activity in the bulb, we examined the mathematical structure of the maximum entropy distribution, the **h_i_** and **J_ij_** terms. The local field term **h_i_**, was always significantly higher during running as compared to when the animal was stationary (for 11-unit subpopulation: **h_i,running_** 5. 20 ± 1.67, **h_i,stationary_** = −5.42 ± 1.81, p < 0.005, two-sided Wilcoxon signed-rank test, **Fig. 5E**, right) across subpopulations ranging in size from 3 to 19 neurons (p < 10^-6^, two-sided Wilcoxon signed-rank test with Bonferroni correction, **Fig. 5F**), consistent with the mean increase in firing rates observed when the animal ran (**Fig. 2D**). Interestingly, the distributions of the **J_ij_** terms which captured the interactions between neurons were significantly decreased during running (for 11-unit subpopulation: **J_ij,stationary_** = 0.41 ± 1.15; **J_ij,running_** = −0.08 ± 0.95, p < 0.005, two-sided Wilcoxon signed-rank test, **Fig. 5G**, right). Furthermore, while the mean **J_ij_** term was positive during stationary epochs across an array of subpopulation sizes (3-19 neurons), the **J_ij_** terms were slightly negative during running epochs, (p < 10^-6^ for all population sizes, two-sided Wilcoxon signed-rank test with Bonferroni correction, **Fig. 5H**). This result, combined with our entropy findings (**Fig. 4A**), provides insight into the poorer fit of the model during running epochs. While the increased entropy showed that population patterns in the MOB became more diverse with running (**Fig. 4**), the diminished **J_ij_** interaction terms during running meant the pair-wise maximum entropy models were comparatively insufficient to capture the increased complexity of ensemble activity.

Previous work has suggested that the interaction term **J_ij_** is related to pairwise correlations between neurons (Schneidman et al. 2006). To explore this, we calculated the pair-wise correlation for every pair of units in each animal during stationary and running epochs (**Fig. 6A**). While the **J_ij_** terms were estimated and then iteratively updated to improve their fit (see *Materials and Methods)* (Maoz and Schneidman 2017), the pairwise correlation was directly calculated from continuous firing rate time-series. They thus represented two complementary methods for assessing pairwise interactions. We found that the pairwise correlations during running were smaller than that during stationary epochs (**Fig. 6B,C**), consistent with the finding of **J_ij_** terms based on behavior (**Fig. 5G,H**), demonstrating that during locomotion, not only the firing rates but the interactions between neurons are affected.

**Figure 6:**
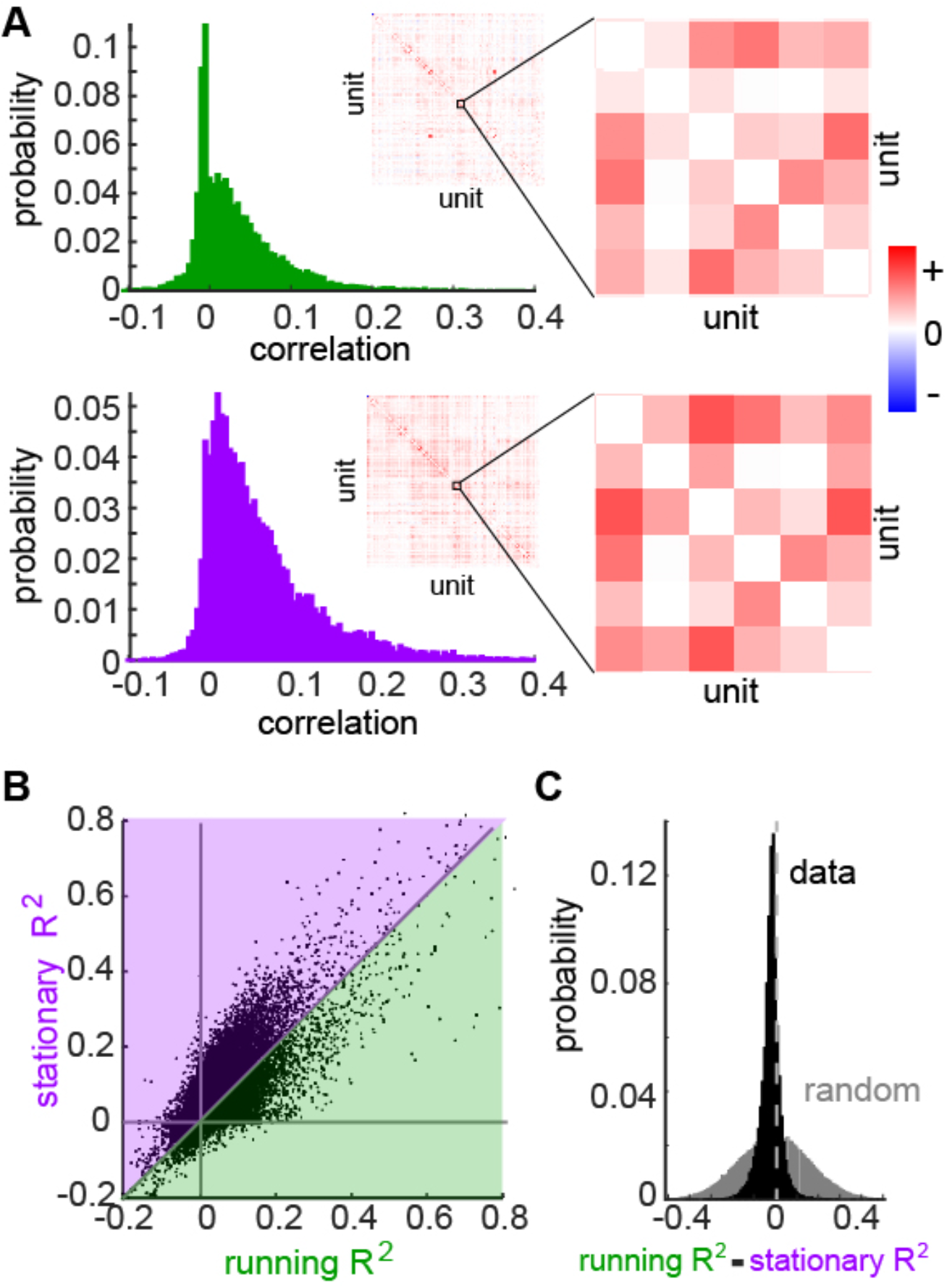
M/T cell pair-wise correlations were lower when animal was running. (A) Histogram of M/T cell pair-wise correlations for a single animal when the animal was running (top) versus when the animal was stationary (bottom). (A, insert) Correlation matrix for all pairs (positive correlations red, negative correlations =blue). (B) Correlated activity during running plotted against correlated activity when the animal was stationary. (C) Histogram of differences in correlation between when the animal was running versus stationary (black) as compared to shuffled data (gray).

## Discussion

Here we found that mitral/tufted cell population activity in the main olfactory bulb of the mouse changed during the animal’s locomotion. In addition to increases in firing rates of individual neurons, the entropy of population activity was increased when the animal ran, resulting in an increase in the dimensionality of the ensemble activity. A second order maximum entropy model fit to the population activity revealed that weak-pairwise interactions could explain the global structure of the firing when the animal was stationary, but the explanatory power of these pairwise interactions diminished when the animal ran. We therefore suggest that running is associated with higher-order interactions between M/T cells.

What do these changes in neuronal population activity signify in terms of olfactory function? Without recordings of population responses to arrays of odors while the animal is running versus when it remains stationary, the information content of the neuronal ensemble cannot be extrapolated from the entropy. These information estimates depend heavily on how the stimulus domain is partitioned (how different are the odors, odor classes, concentrations, etc.) as well as the sampling (number of stimuli presented, number of trials, etc.) (Hallem and Carlson 2006; Meister and Bonhoeffer 2001; Rubin and Katz 1999; Saito et al. 2009; Soucy et al. 2009), questions we leave for future work. We instead focus on the question of how locomotor behavior reshapes the structure of spontaneous activity in the olfactory bulb. Previous work has shown that running is associated with increased whisking, sniffing, and spontaneous firing rates in M/T cells. Interestingly, while we find this to be true for a number of M/T cells, we also find a number of neurons whose firing rate reduces with running. Additionally, while we were not able to measure sniffing in these experiments, recent studies demonstrate the sniffing in head fixed animals may be different than freely behaving animals (Nguyen Chi et al. 2016). While sniffing, whisking, neural circuits could give rise to changes in both single cell and population M/T cell activity (Carey and Wachowiak 2011; Kleinfeld et al. 2014; Moore et al. 2013; Wachowiak 2011), here we focus on *how* these behaviors during running could influence neural population activity, and what this might mean for general principles of encoding.

First, studies in visual cortex (MacLean et al. 2005; Niell and Stryker 2010), auditory cortex (Bender et al. 2016; Luczak et al. 2009), and the olfactory bulb (Stakic et al. 2011; Thompson et al. 2018) suggest that patterns of spontaneous activity recapitulate patterns of activity in response to sensory stimuli (Carrillo-Reid et al. 2019; MacLean et al. 2005; Tsodyks et al. 1999), effectively outlining the realm of possible patterns that *can* occur in response to inputs (Luczak et al. 2009; Thompson et al. 2018). As entropy increases and pair-wise correlations decrease during locomotion, our results suggest that the realm of possible patterns of activity when the animal was running are larger. Consequently, if spontaneous activity defines a manifold on which responses to odors reside (Chen and Padmanabhan 2020; Thompson et al. 2018), then our study shows that the size of that manifold is larger when the animal runs. Recent evidence suggests that spontaneous activity in the olfactory cortex (piriform cortex) is driven by the firing of neurons in the bulb (Tantirigama et al. 2017). Changes in the structure of spontaneous activity in the bulb during locomotion may therefore dynamically gate the range of olfactory coding in the piriform cortex. Furthermore, as pair-wise correlations are better at describing the patterns that occur when the animal is stationary as compared to when it is running, our work suggests that the space in which olfactory responses reside during locomotion may be more complex, and defined by higher order interactions among neurons (Ohiorhenuan et al. 2010; Schneidman et al. 2011; Tkačik et al. 2014).

How do these functional differences arise? Maximum entropy models can be thought of as a way to generate null hypotheses about the functional connectivity within neural circuits (Schneidman 2016). In the retina for instance, pairwise maximum entropy models are sufficient to account for the joint activity of ensembles of neurons (Schneidman et al. 2006; Shlens et al. 2006), a statistical description of observed patterns that arise from the anatomical constraints of the circuits (Pillow et al. 2008; Shlens et al. 2006). By contrast, these same pair-wise interaction models fail to account for the joint activity of populations as small as 5 neurons in the primary visual cortex (Ohiorhenuan et al. 2010) suggesting that higher order structure (3^rd^, 4^th^ order), possibly arising from interactions with local inhibitory interneurons, may govern the structure of neuronal population activity. In the bulb, the dense interconnectedness between M/T cells and granule cells (Kapoor and Urban 2006; Willhite et al. 2006) suggests that the earliest stages of olfactory processing may be more like cortex than retina. Importantly, although organized in some columnar structure (Willhite et al. 2006), the M/T cell interactions with inhibitory granule cells can extend across large spatial scales throughout the bulb (Burton 2017; Gilra and Bhalla 2015; Migliore et al. 2010), possibly more akin to long range connections in cortical circuits (Ruthazer and Stryker 1996). Additionally, granule cells receive centrifugal inputs from a number of regions including the anterior olfactory nucleus (AON), the piriform cortex, the horizontal diagonal band, and the raphe nucleus (Petzold et al. 2009; Price and Powell 1970; Rothermel and Wachowiak 2014; Shipley and Adamek 1984). Among these, monosynaptic projections from both the ventral CA1 (vCA1) region of the hippocampus and the entorhinal cortex (Davis and Macrides 1981; Padmanabhan et al. 2018) may be candidate circuits relaying locomotion information to the bulb, altering the activity of M/T cells via the local inhibitory interneuron population. We caution that how locomotion is operationalized in this work is experimentally constrained, as animals are head-fixed and run on a cylinder, with behavior delineated into epochs of stationary versus running. Nonetheless, these experimental distinctions between “stationary” and “running” may correspond to ethologically different behavioral modes. Locomotion for instance, is necessarily engaged in odor navigation and tracking (Gire et al. 2016; Khan et al. 2012), while animals are often stationary when sniffing a single object in service of odor discrimination or odor identification (Kepecs et al. 2007; Rinberg et al. 2006b; Uchida and Mainen 2003). These different behavioral modes may require either different underlying coding strategies or may include information about behavior itself in the code relayed by M/T cells. Our results suggest that the structure of spontaneous activity in the bulb is highly flexible to support these different modes, operating within a dynamic high-dimensional space shaped by the animal’s behavior (Tsuno and Mori 2009).

## Authors Contributions

K.P. conceived and supervised the project. K.P. and U.C. performed the analysis, made the figures, and wrote the manuscript. E.J.W.C. and K.P. performed the experiments. R.C.G, S.W. and E.L. assisted with analyses. All authors approved the final version of the manuscript.

## Acknowledgements

KP was supported by NSF CAREER (1749772), NIMH (R01MH11392), the Schmitt Foundation, and the Cystinosis Research Foundation. RCG was supported by NINDS (U19NS112953) and NIDCD (R01DC018455).

